# Synergism of Cry1Aa protein against *Grapholita molesta* (Busck) by a cadherin fragment from *Spodoptera exigua* (Hübner)

**DOI:** 10.64898/2026.07.29.741540

**Authors:** Ascensión Andrés-Garrido, Ayda Khorramnejad, Rosa M. González-Martínez, Baltasar Escriche

## Abstract

*Bacillus thuringiensis* (Bt) is currently the most commercialized biopesticide worldwide. Despite the success of Bt application in biological control programs, diverse strategies have been developed to enhance the toxicity of Bt toxins and delay the emergence of insect resistance, including the addition of cadherin fragments as a synergistic agent. In this study, the synergistic effect of two different cadherin fragments, one from a lepidopteran species, *Spodoptera exigua* (rSeCad1bp), and the other from a coleopteran species, *Tenebrio molitor* (rTmCad1p), was evaluated on the toxicity of Cry1Aa, Cry1Ab, and Cry1Ia against *S. exigua* and *Grapholita molesta*. Our results show that while the toxicity of Cry1 proteins to *S. exigua* was not affected by rSeCad1bp, the toxicity of Cry1Aa against *G. molesta* increased about 2.6-fold. No other synergistic or antagonist effects were observed. The potential mechanisms for the detected toxicity enhancement in *G. molesta* were studied, including protection against proteolysis, promotion of oligomerization, and increased binding to receptors on midgut brush border membrane vesicles (BBMV). Results revealed a slight increase in Cry1Aa binding to BBMV. These data highlight the great potential of using rSeCad1bp in combination with Cry1Aa for control of *G. molesta*.

**Key Contribution:** Our results show that the cadherin fragment from *Spodoptera exigua* synergized Cry1Aa toxicity against *Grapholita molesta*, but not to *S. exigua*. The synergistic effect observed might be partly associated with an increase in Cry1Aa binding to the target midgut membrane.

## 1. Introduction

*Bacillus thuringiensis* (Bt) is a gram-positive bacterium producing different proteins mostly active against insects. Among Bt toxins, vegetative insecticidal proteins (Vip) and crystal proteins (Cry) are mostly used in insect control both in sprayable formulations and transgenic crops [1, 2]. Cry proteins are toxic to species of Lepidoptera, Diptera, Coleoptera, Hymenoptera, Homoptera, Dictyoptera, Orthoptera, and Mallophaga, in addition to nematodes (Strongylida, Tylenchida), protozoa (Diplomonadida), and mites (Acari) [3, 4, 5]. The safety and effectiveness of Bt-based bioinsecticides for the control of many agricultural pests have made them the most successful commercialized bioinsecticides worldwide [6, 7]. However, the emergence of insect pest resistance to both synthetic and biological pesticides is one of the main concerns in agriculture [8].

Management of resistance requires an understanding of the mechanisms involved to identify tools delaying and overcoming resistance [9,10, 11]. Over the years, different strategies were proposed to delay insect resistance to Bt pesticides (i.e., using mixtures of Cry proteins with a different mode of action, employing modified Cry toxins), until Chen et al. [12] proposed a new approach based on the possibility of using cadherin-based Cry toxin enhancers. Cadherins are calcium-dependent transmembrane glycoproteins described as functional receptors for Cry toxins, especially for Cry1A in Lepidoptera [13–18], but also in Coleoptera [19, 20] and Diptera orders [21, 22, 23]. Binding of Cry proteins to cadherin has been revealed as a common step in different models proposed for the mode of action of Cry proteins [5, 24–28]. The domain structure of cadherin receptors usually comprises an internal cytoplasmic domain, a transmembrane region, and an ectodomain formed by a signal peptide, 8-12 cadherin repeats, and a membrane-proximal extracellular domain [29, 30]. The cadherin toxin-binding region usually localizes to the membrane proximal extracellular domain. Several reports have pointed out the importance of cadherin peptides containing the toxin binding region as potential synergists of Cry toxins [12, 20, 23, 31]. This Cry synergistic effect of cadherin fragments has been reported in lepidopteran [12, 32, 33], coleopteran [20, 34], and dipteran insects [23, 35]. For instance, the synergistic effect of cadherin fragments from *Spodoptera exigua* (rSeCad1bp) and *Tenebrio molitor* (rTmCad1p) when co-applied with Cry1Ac, Cry1Ca, or Cry3 has been previously reported in different lepidopteran and coleopteran pests [34, 36, 37].

In this study, we tested the potential use of two cadherin fragments to expand the activity range and increase toxicity of Cry1Aa, Cry1Ab, and Cry1Ia toxins against two lepidopteran species, *S. exigua* (Lep.: Noctuidae) and *Grapholita molesta* (Lep.: Tortricidae). A widely distributed polyphagous pest, *S. exigua* larvae feed on more than 20 different plant families and is considered one of the most important pests of field and greenhouse crops. Importantly, the long-term application of synthetic [38] and Bt-based pesticides [11, 39, 40, 41] has led to the development of *S. exigua* resistance, making it essential to develop new effective biopesticides against this pest. Larvae of *G. molesta* are a major pest of stone and pome fruits, such as peach and apple. Currently, pheromonal and chemical controls are the most common tools used to reduce damage of this pest [42, 43]. It is noteworthy to mention that *S. exigua* exhibits low susceptibility to Cry1Aa, Cry1Ab, and Cry1Ia toxins [44, 45]; while *G. molesta* has high susceptibility to Cry1 toxins [46]. The two cadherin fragments used for this study include peptides from *S. exigua* (rSeCad1bp) and *T. molitor* (rTmCad1p) cadherin. rSeCad1bp contains CR7, CR8, CR9, CR10, CR11, and MPED regions, while rTmCad1p contains CR12 and MPED regions. Combinations of these cadherin peptides with Cry1Aa, Cry1Ab, and Cry1Ia were tested to explore their possible synergistic activity against *S. exigua* and *G. molesta*. Notably, Cry1Ia shares less than 70% similarity with Cry1Aa and Cry1Ab, while both Cry1A proteins share more than 95% sequence similarity. Our results support that rSeCad1bp synergistically increases the toxicity of Cry1Aa against *G. molesta*, the first report of a cadherin-Cry toxin synergism in *G. molesta*. Based on data reported previously, experiments were designed to identify the molecular bases for this synergism by analyzing differences in steps in the Bt-mode-of-action: Cry toxins stability in the midgut [47]; formation of cadherin-induced toxin oligomers [19, 31, 33], and Cry toxins ability to bind to midgut brush border [12, 35].

## 2. Results

### 2.1. rSeCad1bp enhanced Cry1Aa toxicity against *G. molesta*

To test the effectiveness and accurate preparation of Cry toxins, the susceptibility of first-instar *G. molesta* and *S. exigua* larvae to Cry1Aa, Cry1Ab, and Cry1Ia proteins was preliminarily determined by performing single-dose bioassays at a toxin concentration of 1,000 ng/cm^2^ (Figure 1).

**Figure 1.**
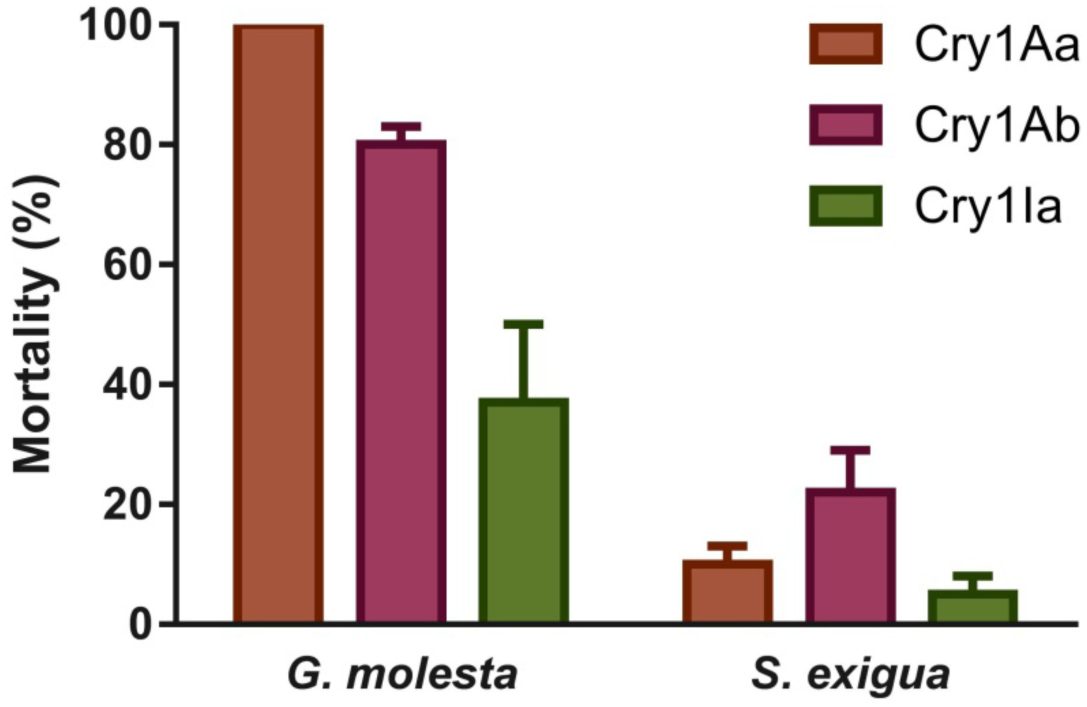
Susceptibility of *G. molesta* and *S. exigua* to Cry toxins at 1,000 ng/cm^2^. The mean and the standard error of the mean (SEM) are represented.

The bioassay results revealed that Cry1Aa and Cry1Ab proteins were highly toxic to *G. molesta* (100% and 80% mortality, respectively), while Cry1Ia caused about 40% mortality. In contrast, all the toxins were marginally toxic to *S. exigua* since mortality values were lower than 25% in all experiments.

To determine whether the addition of rSeCad1bp or rTmCad1p fragments leads to increased Cry1 toxicity, single-dose bioassays were carried out using several toxin: cadherin fragment molar ratios (1:0.1, 1:1, and 1:10) (Figure 2). The Cry1A proteins alone yielded about 35% mortality for *G. molesta*, while less than 20% mortality was observed for Cry1Ia, as with *S. exigua* larvae. Fragments of cadherin alone (control) were not toxic to either insect. In contrast, a significant ∼2-fold toxicity enhancement against *G. molesta* was observed with rSeCad1bp:Cry1Aa mixtures (Figure 2a). The addition of rSeCad1bp to Cry1Aa at the 1:0.1, 1:1, and 1:10 ratios resulted in a significant increase in *G. molesta* larval mortality from approximately 37% to 75%, 88%, and 94%, respectively. The differences in mortality obtained among these ratios were not statistically significant. On the other hand, mixtures of rSeCad1bp:Cry1Ab seemed to result in higher toxicity compared to Cry1Ab alone, but differences were not statistically significant. Similarly, while rTmCad1p:Cry1Aa mixtures resulted in higher toxicity compared to Cry1Aa alone, only increased mortality at the 1:1 ratio, with mortality increasing from 39% to 78% (Figure 2b), was statistically significant. No significant effects were detected for mixtures in *S. exigua* larvae, except for Cry1Ia: rSeCad1bp (1:1) (Figures 2c-2d). Although academically interesting, further exploration of Cry1Ia synergism was discarded due to the low mortality achieved, which complicated pest control and commercial interest in it.

**Figure 2.**
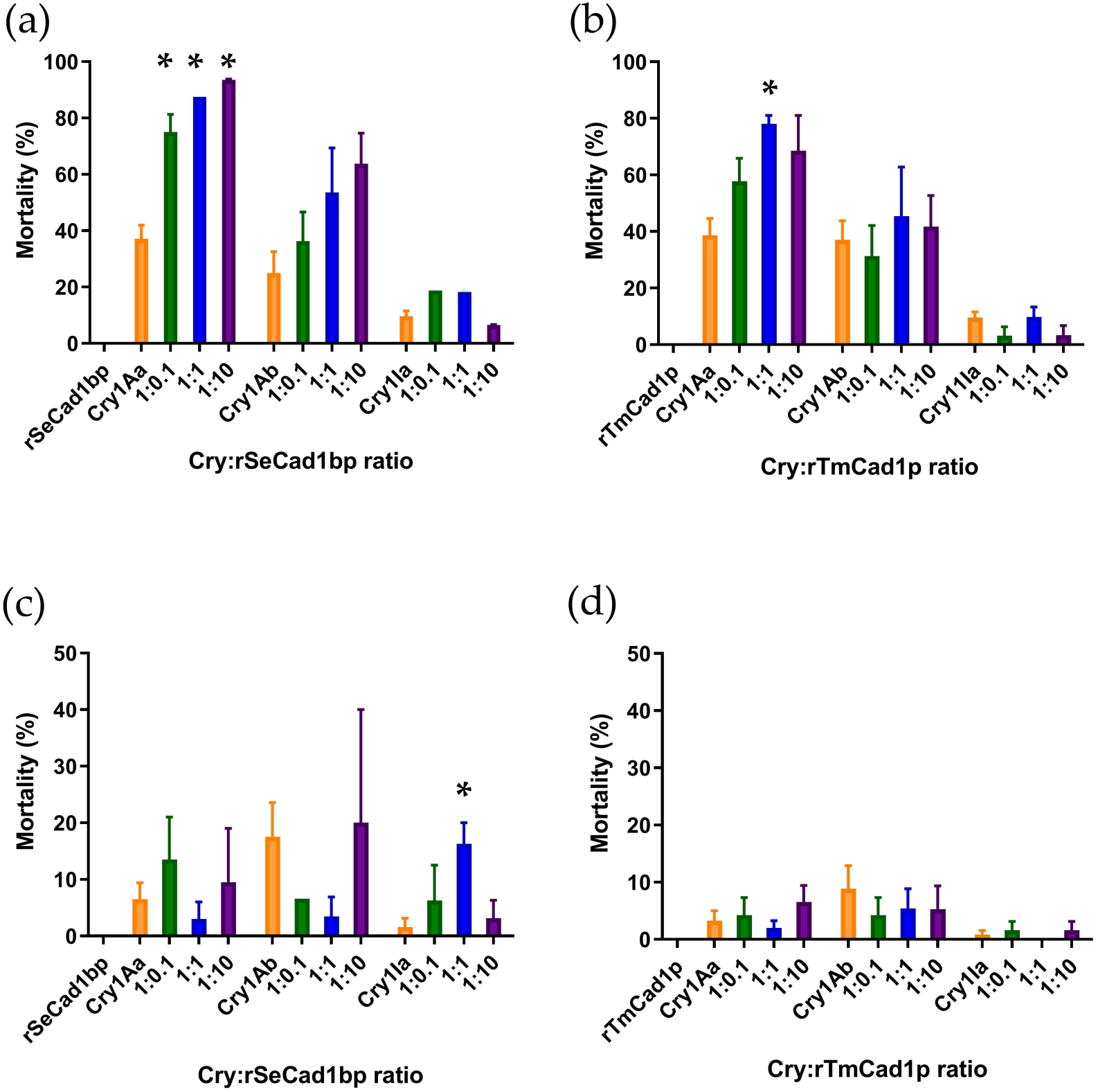
Toxicity of Cry1 and cadherin fragments. alone or in combination, on *G. molesta* (a and b) and *S. exigua* (c and d) larvae. Cry1 toxins (10 ng/cm2) were used at various molar ratios of rSeCad1bp (a and c) or rTmCad1p (b and d). The selected concentration of Cry1 toxins was based on practical and economical application. The mean and t the standard error of the mean (SEM) are represented. Bars are not shown where mortality was 0% after treatment. Significant differences (one-way ANOVA with post hoc analysis, p < 0.05) in mean percentage mortality, compared to Cry1 alone, are indicated by an asterisk above the bars.

Dose-response bioassays were designed to more exactly determine the synergistic effect of cadherin fragments and Cry1Aa against *G. molesta* (Table 1). The influence of rSeCad1bp on Cry1Aa toxicity toward *G. molesta* at the ratio of 1:0.1 was chosen for further analysis based on the practical application principle that the synergistic factor should be present in a lower amount than the active ingredient.

**Table 1.** Parameters of the dose-response bioassays on *G. molesta* challenged with Cry1Aa alone or in combination with rSeCad1bp and rTmCad1p cadherin fragments. Toxin concentration ranged from 0 to 270 ng/cm^2^. Probit analysis provided the values for LC_50_ with its 95% fiducial limits (FL_95_) and slope with its standard deviation (SD).

| Protein | No. of larvae | LC<sub>50</sub> (ng/cm<sup>2</sup>) | FL<sub>95</sub><br>(Lower-upper) | Slope ± SD |
| --- | --- | --- | --- | --- |
| Cry1Aa | 446 | 9.9 | 5.8-15 | 2.21 ± 0.27 |
| Cry1Aa: rSeCad1bp<br>(1:0.1) | 220 | 3.7 | 2.5-5.2 | 1.77 ± 0.23 |
| Cry1Aa: rTmCad1p<br>(1:1) | 217 | 6.6 | 2.7-11.2 | 2.46 ± 0.4 |

In agreement with the results obtained in the single-dose assay, the LC_50_ value obtained with rSeCad1bp addition (3.7 ng/cm^2^) was lower (more toxic) than that of Cry1Aa alone (9.9 ng/cm^2^), which means that less amount of Cry1Aa is necessary to reach 50% of larval mortality. Assays with and without rSeCad1bp produced parallel curves and slopes which were not significantly different, suggesting that the synergism is not only found at a specific concentration (i.e. 10 ng/cm^2^) but at a range of concentrations (from 0 to 270 ng/cm^2^). Thus, the potency of the mixture was calculated [48]. Probit analyses estimated a 2.6 (FL_95_: 1.5-4.2) fold increase in Cry1Aa toxicity for the 1:0.1 mixture with rSeCad1bp. The experiments with Cry1Aa mixed with rTmCad1p provided no significant differences when compared to Cry1Aa alone (Probit analysis), indicating that significant toxicity enhancement was not achieved at any Cry1Aa concentration.

### 2.2. rSeCad1bp and rTmCad1p interact with Cry1Aa toxin

Dot blot assays were performed to test the interaction of Cry1Aa protein with cadherin fragments (Figure 3). The results showed that Cry1Aa protein was able to interact with both rSeCad1bp and rTmCad1p fragments in a dose-dependent manner. Qualitative comparison between both peptides showed that the signal obtained for rTmCad1p was stronger than rSeCad1bp at all the concentrations tested, which may be attributed to differences in cadherin fragment size since rTmCad1p was smaller than rSeCad1bp for the same amount employed, i.e. 2 µg, the Cry: cadherin molar ratio changes). These results suggested direct interaction between the cadherin fragments (rSeCad1bp and rTmCad1p) and Cry1Aa.

**Figure 3.**
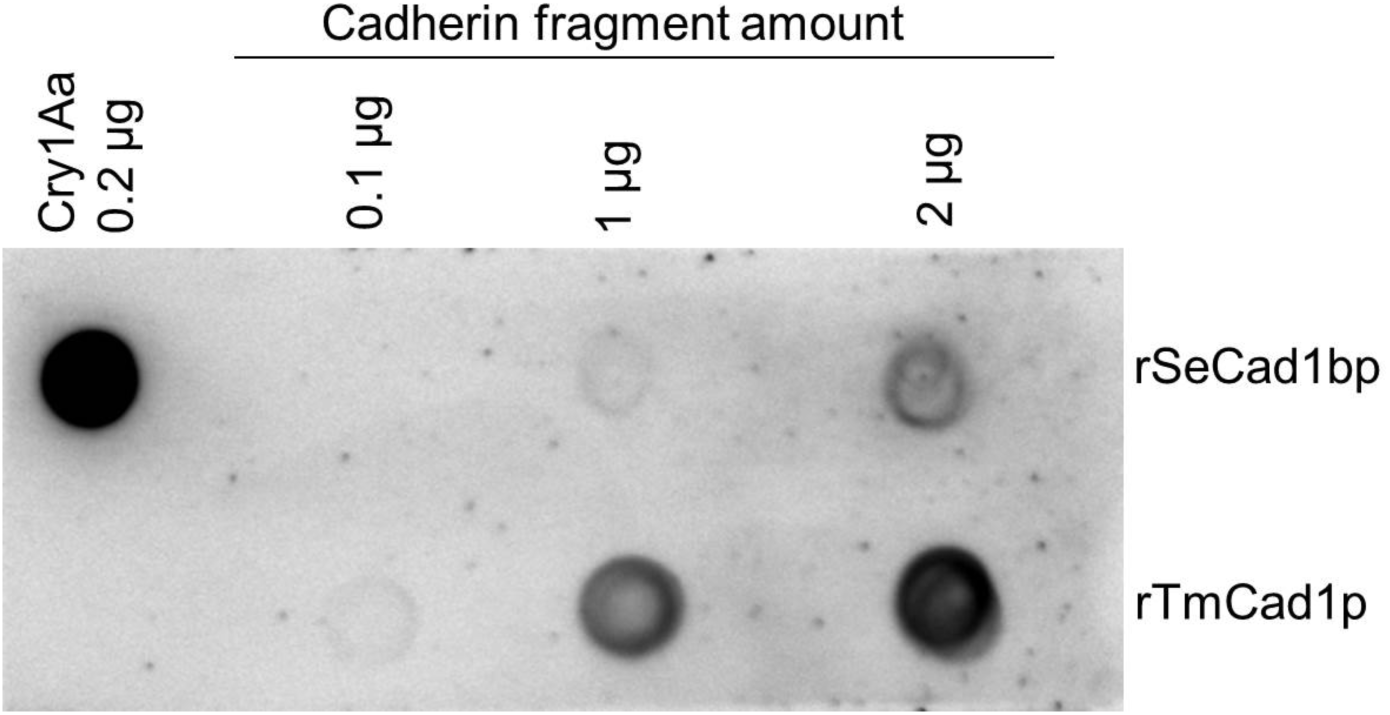
Dot blot assay of rSeCad1bp and rTmCad1p interaction with Cry1Aa. Increasing amounts of cadherin fragments were spotted on the PVDF membrane, and after blocking, probed with 20 ng/ml of Cry1Aa. Bound Cry1Aa was detected with anti-Cry1A polyclonal serum followed by secondary antibody. Cry1Aa (200 ng) spotted alone was used as a positive control.

### 2.3. rSeCad1bp does not prolong the stability of Cry1Aa in the presence of *G. molesta* midgut fluids

In general, Cry toxins are processed and activated by the action of proteases in the midgut fluids. However, in some cases these proteases can also degrade the Cry protein, thus disrupting receptor-binding and pore-forming ability [49, 50]. One of the bases of the synergistic effect of the cadherin fragments has been attributed to an increase in toxin stability to these midgut proteases [47]. We tested whether interactions between rSeCad1bp and Cry1Aa increased the stability of Cry1Aa in the presence of midgut fluid proteases, which could explain the observed synergism in *G. molesta*. The protection of Cry1Aa toxin by the rSeCad1bp cadherin fragment was tested *in vitro* (Figure 4).

**Figure 4.**
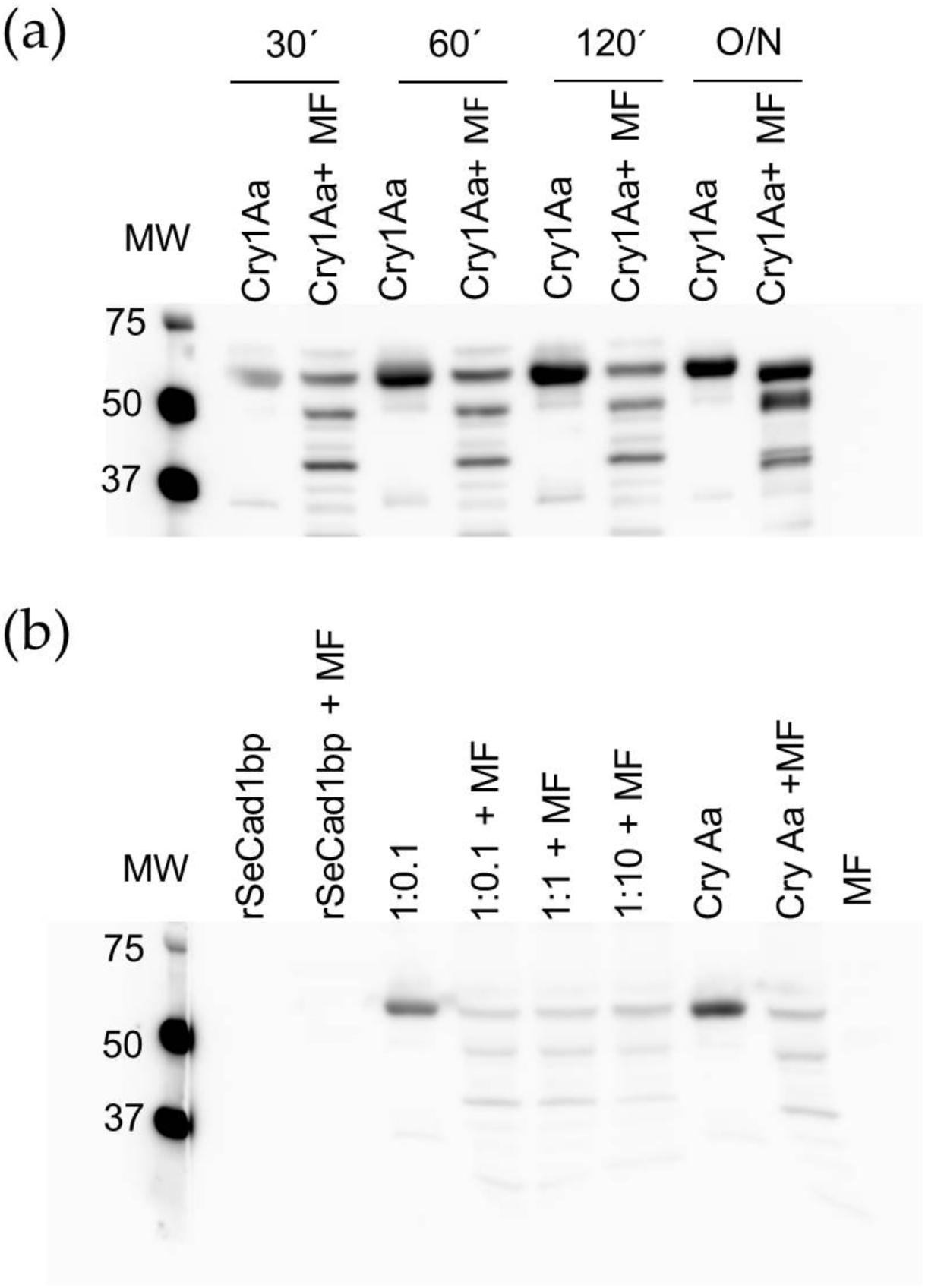
Stability of Cry1Aa processing by *G. molesta* midgut fluids (MF) in the presence or absence of rSeCad1bp. (a*) In vitro* time course of proteolytic processing of Cry1Aa. (b) Analysis of Cry1Aa degradation protection by rSecad1bp after incubation with *G. molesta* MF for 30 min. MW, Precision Plus Protein™ Dual Color molecular mass marker (in kDa) (Bio-Rad, Hercules, CA, USA). MF, proteins present in the midgut fluids from G. molesta. Cry1Aa:rSeCad1bp molar ratios were indicated as 1:0.1, 1:1, and 1:10.

Time course assays (Figure 4a) revealed the processing of Cry1Aa incubated with 10% (vol/vol) of *G. molesta* MF at tested time intervals (30, 60, 120 min, and overnight). Similar band patterns were obtained for all incubation times, suggesting that the proteolysis was produced in less than 30 min. The strong bands observed after overnight incubation could correspond to now solubilized fragments, which precipitated at shorter incubation periods. The absence or presence of rSeCad1bp at several ratios did not change the Cry1Aa band pattern obtained after MF treatment (Figure 4b), suggesting that rSeCad1bp did not increase the stability of Cry1Aa in *G. molesta* midgut fluids.

### 2.4. Toxin-oligomer formation is not responsible for rSeCad1bp-mediated Cry1Aa toxicity enhancement

An alternative molecular hypothesis tested to explain the observed synergistic activity was the potential for increased Cry1 oligomerization by addition of a cadherin fragment. The impact of rSeCad1bp in Cry1Aa oligomerization was studied by Western blotting analyses (Figure 5).

**Figure 5.**
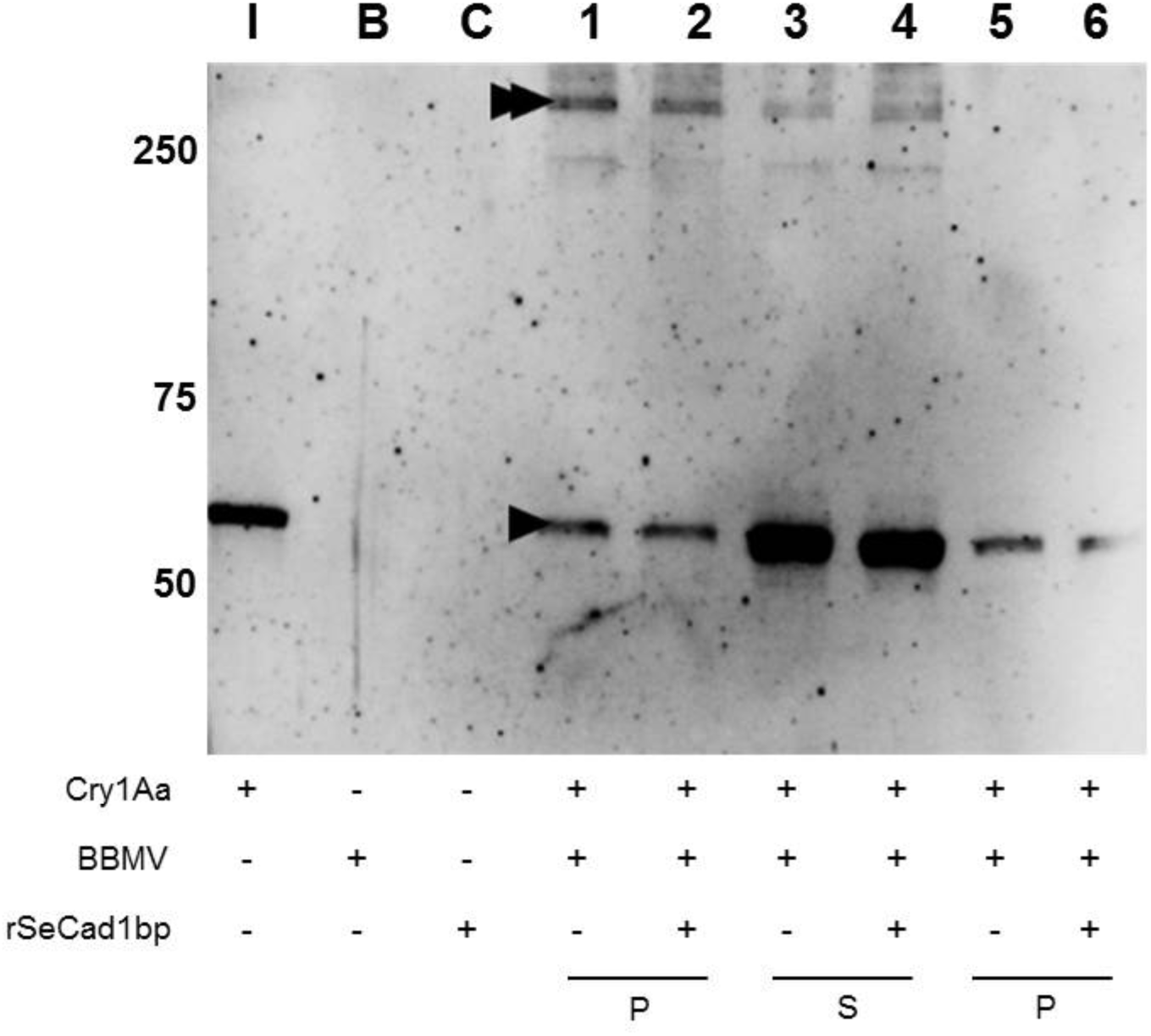
Promotion of the oligomeric structure of Cry1Aa after incubation with *G. molesta* BBMV and rSeCad1bp fragment. Controls include I= Cry1Aa toxin (50 ng), B= *G. molesta* BBMV (10 µg), and C= rSeCad1bp fragment (200 ng). P= Pellet containing Cry1Aa toxin bound to BBMV (lane 1), or Cry1Aa toxin incubated with BBMV and rSeCad1bp fragment (lane 2). S= Supernatant obtained after incubation of Cry1Aa toxin with BBMV (lane 3) or after incubation of Cry1Aa toxin with BBMV and rSeCad1bp fragment (lane 4). Lanes 5 and 6 show Cry1Aa toxin bound to the BBMV heated at 99°C to denature the oligomer (lane 5) and Cry1Aa toxin bound to the BBMV in the presence of rSeCad1bp and heated at 99°C (lane 6). The size of protein monomers (arrowhead) and oligomers (double arrowhead) was estimated using Precision Plus Protein™ Dual Color molecular mass marker (Bio-Rad, Hercules, CA, USA).

The results of the oligomerization assays showed that Cry1Aa was incorporated into *G. molesta* BBMV in both monomeric and oligomeric forms (∼60 kDa for monomer and ∼250 kDa for oligomer) (Figure 5, lane 1). The band size corresponding to the oligomeric structure suggested a tetrameric form, which was denatured after 10 min of heating at 99 °C (Figure 5, lanes 5 and 6). The monomeric and oligomeric Cry1Aa structures were also observed in the supernatant, not bound to the BBMV (Figure 5, lanes 3 and 4). The addition of rSeCad1bp did not have relevant effects on the formation of Cry1Aa oligomers (Figure 5, lane 2).

### 2.5. rSeCad1bp increases binding of Cry1Aa to *G. molesta* BBMV

The effect of rSeCad1bp on the binding of Cry1Aa protein to *G. molesta* BBMV was tested using radiolabeled Cry1Aa (Figure 6).

**Figure 6.**
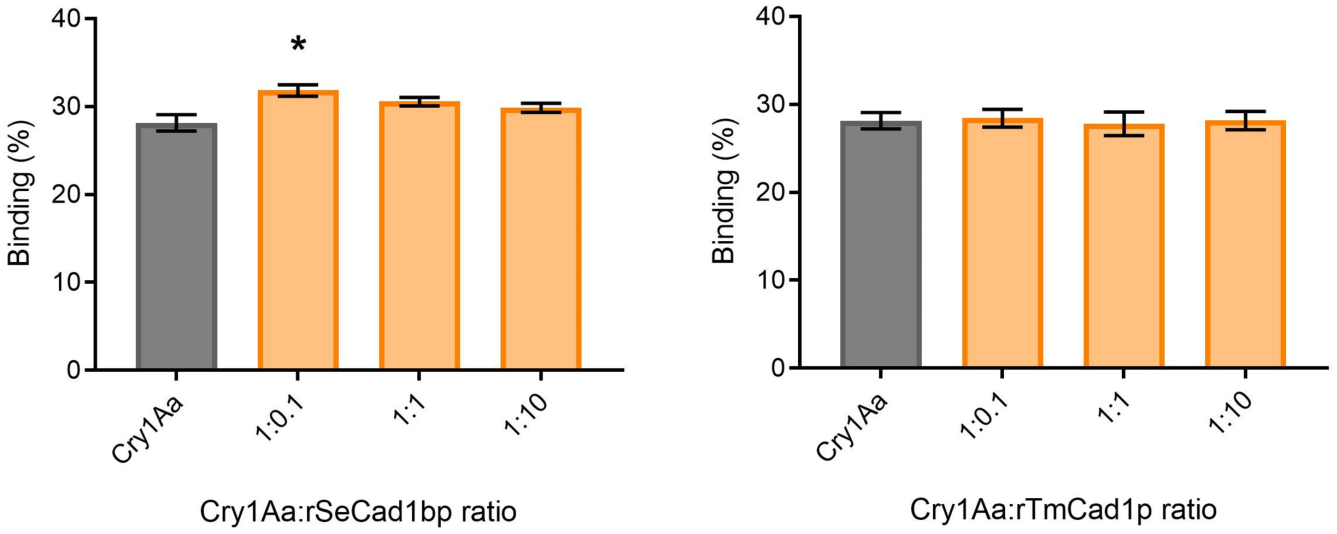
Binding of ^125^I-Cry1Aa to *G. molesta* BBMV in the presence of rSeCad1bp and rTmCad1p. Effect of different molar ratios of rSeCad1bp and rTmCad1p on the total binding of ^125^I-Cry1Aa to *G. molesta* BBMV. An asterisk above the columns indicates significant differences (ANOVA one-way with post-test analysis, p-value < 0.05) between Cry1Aa: cadherin ratios and Cry1Aa alone (as control).

The binding results showed that Cry1Aa binding to *G. molesta* BBMV was only significantly increased when rSeCad1bp was employed at the ratio 1:0.1 (Figure 6). In Figure 6, it can be observed that Cry1Aa binding to 0.15 mg/ml of BBMV is around 28 %, which is consistent with the total binding curve. Furthermore, neither rSeCad1bp nor rTmCad1p blocked Cry1Aa binding to BBMV at the ratios tested.

## 3. Discussion

We studied the synergistic effect of two cadherin fragments from different insect species (rSeCad1bp from *S. exigua* and rTmCad1p from *T. molitor*) on Cry1Aa, Cry1Ab, and Cry1Ia toxicity toward *S. exigua* and *G. molesta*. No antagonistic effect of cadherin fragments on Cry toxicity was found, in contrast with previously reported results [51, 52, 53]. Preliminarily, the enhancement of Cry1 toxicity by rSeCad1bp and rTmCad1p was assessed in single-concentration experiments. We found that the susceptibility of *G. molesta* and *S. exigua* to Cry1Aa, Cry1Ab, and Cry1Ia was in agreement with previous studies [44–46, 54], except for the newly described toxicity of Cry1Ia to *G. molesta*. In our bioassays, significant enhancement of Cry1 toxicity by cadherin fragments was only detected for rSeCad1bp:Cry1Aa and rTmCad1p:Cry1Aa mixtures toward *G. molesta*. The toxicity enhancement by rSeCad1bp was found at all the ratios tested, while rTmCad1p synergistic effect was only observed at the 1:1 molar ratio. These results are in agreement with previous studies showing enhancement of Cry1A toxicity by cadherin fragments [12, 31, 33, 55].

The lack of synergism in *S. exigua* differed from the previously reported role of rSeCad1bp as a synergist of Cry1B and Cry1C toxicity in *S. exigua* [32,37]. Gao et al. [56] reported that midgut juice could degrade cadherin fragments, especially in low proportions. Since our study used up to a 1:10 ratio compared to the 1:50/1:100 ratio used in previous studies [32, 37], it can be hypothesized that the amount of rSeCad1bp used may account for the difference. In the same way, the absence of a general synergistic effect in *S. exigua* compared to *G. molesta* might be due to differential midgut environments. Similarly, while the rTmCad1p was reported to synergize Cry toxins in several coleopteran and lepidopteran species [19, 34, 36], no activity was detected in our bioassays with *S. exigua*. This lack of rTmCad1p synergism was also reported for Cry1C and Vip3A against *S. exigua* [57] and supports that this cadherin fragment may not work effectively in *S. exigua*.

The single-dose assays performed did not address whether the Cry1 synergism is dose-dependent. No statistically significant differences between the ratios were observed, which confirmed that the synergism in *G. molesta* is not toxin-to-peptide–ratio-dependent, in contrast to previous studies on different insect species wherein the magnitude of the enhancement is correlated with the ratio of toxin to peptide used [12, 31]. The interaction responses (synergism, additive, antagonism) are complicated to assess and predict, due to a variety of factors involved and the few studies conducted on this topic [58–60]. For example, our dose-response bioassay results confirmed the ability of rSeCad1bp to synergize Cry1Aa toxicity to *G. molesta* but did not provide support for the rTmCad1p synergism detected in single-dose bioassays. However, direct interactions with Cry1Aa were detected for both rSeCad1bp and rTmCad1p. This result suggests that simple interaction between toxin and cadherin fragment is not a determinant of the observed synergism, as previously reported [23, 35]. For instance, the conformational structure of cadherin peptides could be a variable affecting the relevance of the observed synergistic response. Although it has been suggested that the unfolded structure of cadherin peptides could have more exposed amino acids and result in better Cry interactions [12, 31, 52], in this study both peptides were purified and refolded. While a previous report documented rSeCad1bp synergism of Cry1Ca in *S. exigua* [37], our study is the first report of rSeCad1bp synergizing Cry1Aa in *G. molesta*. These observations support that cadherin synergism of Cry toxicity may occur when using a cadherin derived from a different insect [55], although not in all cases.

The molecular basis of Cry1 synergism by cadherin fragments has been attributed to i) an increase of Cry toxin stability in the midgut [47]; ii) formation of cadherin-induced toxin oligomers [19, 31, 33]; and iii) an increased binding of Cry toxin to the midgut brush border [12, 35, 61]. In all the models proposed for Cry protein mode of action, activation of protoxin to toxin by gut proteases is one of the essential steps [62], and altered proteolytic degradation of the toxin has been associated with resistance [11]. Accordingly, protection of Cry toxin from midgut proteases by a cadherin fragment can lead to an enhancement in Cry toxicity [47]. However, our in vitro proteolytic processing results support that rSeCad1bp had no noticeable effect on Cry1Aa stability under the conditions tested.

The second hypothesis tackled was the possibility of cadherin fragments enhancing Cry1Aa oligomer formation. The Cry1Aa oligomers are expected to represent a tetramer according to assays with lipid bilayers [63]. Oligomers observed in the present study with *G. molesta* BBMV had the same size as those observed previously with *Ephestia kuehniella* BBMV when using a *Bombyx mori* cadherin fragment [64, 65]. However, the addition of rSeCad1bp cadherin did not increase Cry1Aa oligomer formation under the conditions tested, supporting that this is not likely the mechanism for synergism. In fact, the relationship between the formation of cadherin-induced toxin oligomers and synergism is not unidirectional, since the presence of oligomer formation has also been related to a decrease in toxicity [52].

A moderate increase of Cry1Aa binding to *G. molesta* BBMV in the presence of rSeCad1bp was positively evidenced in our assays. The slight increment on binding led us to hypothesize that cadherin action may involve other mechanisms in addition to the three addressed here [66]. Several studies have demonstrated that cadherin fragments reported as synergists could bind to Cry toxins and to BBMV, without blocking Cry toxin binding [12, 31, 37]. It can be speculated that, in our study, *S. exigua* tolerance to Cry1Aa might not be based on the cadherin fragment misfunction, since rSeCad1bp enhances Cry1A toxicity to *G. molesta*. Nowadays, various reports studying the ability of cadherin fragments to enhance Cry protein toxicity are available. It is generally accepted that cadherin fragments can be used as a tool to enhance Cry toxicity. However, the specific combination of cadherin-Cry toxin that will potentiate Cry toxicity toward a specific target pest and the molecular basis of their interaction are still unclear. Particular observations with single concentrations are academically interesting, but obtaining consistent results is necessary to uncover the origin of the synergistic effects. Here, we provide novel data on the absence and presence of synergism among different toxin-cadherin-pest combinations. Furthermore, the ability of rSeCad1bp to increase the toxicity of Cry1Aa against *G. molesta* has been evidenced, and different hypotheses have been discussed and experimentally addressed to unravel the molecular bases of the synergism. In this way, the synergistic effect observed might be partly associated with an increase in Cry1Aa binding to the target midgut membrane. Although further research must be done, our findings underscore the significant potential of combining rSeCad1bp with Cry1Aa to effectively control *G. molesta*.

## 4. Conclusions

The goal of this study was to enhance the insecticidal potency and extend the target range of Cry1Aa, Cry1Ab, and Cry1Ia by employing two different cadherin fragments. Among all the combinations assayed, only the cadherin fragment from *S. exigua* (rSeCad1bp) enhanced 2.6-fold Cry1Aa toxicity against *G. molesta* at the toxin: cadherin molar ratio of 1:0.1. Results suggested that cadherin-mediated enhancement of Cry1Aa toxicity might be associated with an increase in Cry binding to BBMV.

## 5. Materials and Methods

### 5.1. Insect rearing

The strain of *S. exigua* used in this study was a FRA colony kindly supplied by Dr. M. López-Ferber, INRA (St Christol les Alés, France), while the *G. molesta* colony was originally obtained from Entomos AG (Switzerland). Both colonies were established and maintained at the University of Valencia (Spain) facilities, reared on artificial [67] and semi-artificial diet [68], respectively, under controlled conditions of temperature (25 ± 2 °C), humidity (70% RH), and photoperiod (16:8 h light: dark) [69]. The same diet and conditions were used in the bioassays.

### 5.2. Cry proteins preparation

Cry1Aa and Cry1Ab were obtained from recombinant *Escherichia coli* strains provided by Dr. R.A. de Maagd (Wageningen University, The Netherlands). Both proteins were produced following the protocol described by Herrero et al. [70]. Both protoxins were solubilized by incubation overnight in solubilization buffer (50 mM sodium carbonate, 100 mM sodium chloride, pH 10.5) containing 10 mM dithiothreitol. Cry1Ia source and its expression, purification, and solubilization were performed as described previously by Khorramnejad et al. [71]. In order to obtain homogeneous data, activated toxins were used in all experiments. Protoxins activation was performed by adding trypsin at a ratio of 10:1 (protein: trypsin). The activated proteins were visualized by 12% sodium dodecyl sulfate polyacrylamide gel electrophoresis (SDS-PAGE), and their concentration was determined by densitometry using TotalLab™ software (version 12.3, Newcastle, UK), employing bovine serum albumin (BSA) as standard.

For the oligomerization assay, the activated Cry1Aa protein was dialyzed in 20 mM Tris-HCl, 300 mM NaCl (pH 8.5) buffer and purified by anion-exchange chromatography in a HiTrap Q HP column using an ÄKTA 100 Explorer system (GE Healthcare, Amersham, UK) following Crava et al. [72] methodology.

### 5.3. Cadherin fragments preparation

The cadherin fragment from *T. molitor* (rTmCad1p) [34] was generously provided by Dr. J. L. Jurat-Fuentes (University of Tennessee, Knoxville, TN, USA). The cadherin fragment from *S. exigua* (rSeCad1bp) was cloned in pET-30a(+) vector by GenScript Corporation (Piscataway, NJ, USA), optimizing the process by conducting codon optimization for its expression in *E. coli*. Transformation, expression, and purification were performed as described by Ren et al. [37]. The molecular mass of the peptides is ap-proximately 26 kDa (rTmCad1p) and 65 KDa (rSeCad1bp).

### 5.4. Preparation of midgut fluid (MF)

To prepare midgut fluid, midguts of G. molesta fifth-instar larvae were dissected, pooled, and centrifuged at 16,100 x g for 10 min at 4 °C. The supernatant was recovered and stored at -80 °C as detailed by Rahman et al. [47].

### 5.5. Bioassays to assess Cry1 toxicity enhancement by rSeCad1bp and rTmCad1p

*G. molesta* and *S. exigua* bioassays were performed as previously described by Andrés-Garrido et al. [57]. Briefly, 50 µl of the Cry protein sample was spread in each well of bioassay trays. After drying, one neonate was placed in each well and 16 larvae were used per concentration. All bioassays were repeated three times. Mortality was recorded after 7 days. The susceptibility of *G. molesta* and *S. exigua* to Cry1Aa, Cry1Ab, and Cry1Ia protein was tested at a single concentration of 1,000 ng/cm^2^.

The practical and economical synergistic application should be based on a low amount of toxin. Accordingly, the concentration of 10 ng/cm^2^ as a practical low concentration of the toxin was chosen. Cry1Aa/Cry1Ab/Cry1Ia toxins with rSeCad1bp or rTmCad1p at the molar ratios of 1:0.1; 1:1; 1:10 (toxin: cadherin fragment) were applied on the diet surface. Toxin solution buffers and cadherin fragments alone were tested individually as negative controls. Dose-response bioassays using several toxin concentrations with a fixed molar ratio of cadherin fragment were performed to check the concentration range of activity of the synergism. Controls provided less than 10% of mortality in all experiments.

### 5.6. Dot blot assays

Dot blot assays were performed following Peng et al. [31] protocol with some modifications. Several amounts of rSeCad1bp and rTmCad1p (0.1 µg, 1 µg, 2 µg) and Cry1Aa (0.2 µg) were dotted onto a nitrocellulose membrane. After blocking with 5% skimmed milk in phosphate-buffered saline (PBS; 8 mM Na_2_HPO_4_ and 2 mM KH_2_PO_4_, 137 mM NaCl, 2.7 mM KCl, pH 7.4), with 0.1% Tween 20 (PBS-T) overnight at 4 °C, the nitrocellulose membrane was bathed in 20 ng/ml Cry1Aa for 1 hour at room temperature. After incubation, the membrane was washed three times with PBS-T buffer. Anti-Cry1A (1/10,000) polyclonal serum (Abraxis, Warminster, PA, USA) was used as the first antibody for 2 hours, followed by the secondary antibody, anti-rabbit IgG-conjugated horseradish peroxidase (Sigma-Aldrich, Saint Louis, MO, USA) (1/20,000) for 60 min. The results were visualized with SuperSignal West Femto chemiluminescent HRP substrate (Thermo Scientific, Waltham, MA, USA), and an ImageQuant LAS400 image analyzer (GE Healthcare Bio-Sciences, Uppsala, Sweden). Each experiment was repeated at least twice.

### 5.7. *In vitro* protease protection assay

Cry1Aa stability in the midgut was tested by a modification of the method of Rahman et al. [47]. Six µg of the toxin was incubated with MF of *G. molesta* in PBS (10% MF volume/total volume) for 30, 60, 120 min, and overnight at 30 °C. At the end of the incubation time, Cry1Aa proteolysis was stopped by adding Laemmli sample buffer (Bio-Rad, Hercules, CA, USA) and heating the samples at 100°C for 10 min. Samples were separated in 12% SDS-PAGE, transferred onto a PVDF membrane, blocked with skimmed milk (5%), and detected following the same protocol as described for dot blot assays.

To evaluate the effect of rSeCab1bp on Cry1Aa stability in the midgut, Cry1Aa (3 µg) was preincubated with rSeCab1bp for 1 hour at 30 °C at different toxin: rSeCad1bp molar ratios (1:0.1; 1:1; 1:10). After preincubation, Cry1Aa alone, or in the presence of rSeCad1bp, was digested for 30 min at 30 °C with 10% midgut fluid of *G. molesta*. Cry1Aa proteolysis was stopped and detected as previously described to test Cry1Aa stability. Each experiment was repeated at least twice.

### 5.8. Brush border membrane vesicles preparation

The whole last-instar larvae of *G. molesta* were used for brush border membrane vesicles (BBMV) preparation. BBMV were prepared based on the differential magnesium precipitation method [73,74], and the protein concentration of *G. molesta* BBMV was quantified by Bradford [75] using BSA as standard. The prepared BBMV were immediately aliquoted, snap-frozen in liquid nitrogen, and kept at -80 °C until used.

### 5.9. Cry1Aa oligomerization assay

The formation of the oligomeric structure of Cry1Aa protein promoted by *G. molesta* BBMV was studied by Western blot analysis following Khorramnejad et al. [71] with slight modifications. After several preliminary experiments, the oligomerization assays were set up by incubating Cry1Aa (2 µg) with *G. molesta* BBMV (10 µg) in a final volume of 50 µl of 20 mM Tris-HCl buffer (pH 8.5). In order to determine whether rSeCad1bp synergizes with the oligomeric structure of Cry1Aa protein promoted by *G. molesta* BBMV, the rSeCad1bp fragment was added to the reaction in the ratio of 1:0.1 (toxin: rSeCad1bp). The reaction tubes were kept for one hour at 37 °C. The BBMV with bound proteins was separated by centrifugation at 21,100 ×g for 15 min at 4 °C. The supernatant containing the unbound protein was kept for analysis. The pellet was washed with 100 µl of ice-cold 20 mM Tris-HCl buffer (pH 8.5) and centrifuged at 21,100 ×g for 15 min at 4 °C. The final pellet was resuspended in 10 µl of the same buffer, mixed with Laemmli sample buffer, and incubated at 50 °C for 3 min. Due to the sensitivity of the oligomeric structure to high temperatures, the BBMV with bound proteins were also heated at 99 °C for 10 min to dissociate into monomer form to confirm that the observed signal corresponds to the oligomeric status. The proteins were separated by 10% SDS-PAGE, transferred onto PVDF Western blot membrane, and detected with the same protocol as described for dot blot assays except for the use of ECLTM Prime Western blotting detection reagent (GE Healthcare, Little Chalfont, UK). Each experiment was repeated at least three times.

### 5.10. Binding assay

The purified Cry1A protein (25 µg) was labeled with 0.3 mCi of ^125^I (PerkinElmer, Boston, MA, USA) using the Chloramine T method [76]. The specific activity obtained for ^125^I-Cry1Aa was 0.74 µCi/µg. When used for this study, the specific activity had decreased to 0.16 µCi/µg.

*G. molesta* BBMV were centrifuged for 10 min at 16,000 ×g 4°C and suspended in binding buffer (PBS, 0.1% BSA, pH 7.4). To determine the optimal concentration of BBMV to assess the role of cadherin fragments on Cry1A binding, increasing amounts of *G. molesta* BBMV (0-0.4 mg/ml) were incubated with 4.30 nM of ^125^I-Cry1Aa in a final volume of 100 μl of binding buffer for 1 hour at room temperature. An excess of unlabeled protein (150 nM) was used to determine the non-specific binding. After incubation, samples were centrifuged at 16,000 ×g for 10 min, and the pellets were washed twice with 0.5 ml of ice-cold binding buffer. Radioactivity associated with the BBMV was measured in a Gamma counter (2480 WIZARD2 automatic gamma counter, PerkinElmer) and was considered an indication of the Cry1Aa bound. Specific binding was obtained by subtracting the non-specific binding from the total binding observed.

Experiments to study the effect of cadherin fragments on Cry1Aa binding were performed by incubating 0.15 mg/mL of BBMV with ^125^I-labeled proteins. rSeCad1bp or rTmCad1p were added to the reaction in the molar ratio of 1:0.1, 1:1, and 1:10 (toxin: cadherin fragment) in a final volume of 100 µl for one hour at room temperature. Centrifugation, washing, and radioactivity measurement were done as explained above. Binding (%) was calculated by dividing final Cry1Aa total binding by the initial Cry1Aa radioactivity assayed. Each experiment was repeated four times, and assays were performed in parallel.

### 5.11. Statistical analysis

The mean and the standard error of the mean (SEM) are shown and represented unless it was indicated. Mean values were compared by one-way ANOVA multiple comparison test and, if differences were considered significant (p < 0.05), Tukeýs post-test among pairs of treatments was employed to determine their differences, considering significantly different values when p < 0.05. GraphPad Prism version 7.00 software (GraphPad Inc., La Jolla, CA, US) was used for performing the statistical analyses.

LC_50_ values (lethal concentration which produces 50% mortality), 95% fiducial limits (FL_95_), and slope values (change in mortality per unit change in dose) were determined by Probit analysis [77] using the POLO-PC software program (LeOra software, Berkeley, CA, USA) [78]. Comparative potencies were calculated by the program for the experiments with parallel responses in toxicity.

## Author Contributions

Conceptualization, A.A.G. and B.E.; Formal analysis, A.A.G. and B.E.; Funding acquisition, B.E.; Investigation, A.A.G., A.K., R.M.G.M.; Methodology, A.A.G., A.K. and B.E.; Project administration, B.E.; Resources, B.E.; Supervision, B.E.; Visualization, A.A.G. and A. K.; Writing – original draft, A.A.G., A. K. and B.E.; Writing – review & editing, B.E. All authors have read and agreed to the published version of the manuscript.

## Funding

This research was funded by the Spanish Ministry of Science and Innovation (AGL2015-70584-C2-1-R and PID2021-122914OB-I00 (co-funded by EU FEDER funds) and the Generalitat Valenciana grant (CIPROM/2023/56). A.A.G. is a beneficiary of a Spanish Ministry of Science, Innovation, and Universities FPI grant (BES 2016-079134).

## Conflicts of Interest

No potential conflict of interest was reported by the authors.

## Notes

### Competing Interest Statement

The authors have declared no competing interest.

